# Evolution of organismal stoichiometry in a 50,000-generation experiment with *Escherichia coli*

**DOI:** 10.1101/021360

**Authors:** Caroline B. Turner, Brian D. Wade, Justin R. Meyer, Richard E. Lenski

## Abstract

Organismal stoichiometry refers to the relative proportion of chemical elements in the biomass of organisms, and it can have important effects on ecological interactions from population to ecosystem scales. Although stoichiometry has been studied extensively from an ecological perspective, little is known about rates and directions of evolutionary changes in elemental composition in response to nutrient limitation. We measured carbon, nitrogen, and phosphorus content of *Escherichia coli* evolved under controlled carbon-limited conditions for 50,000 generations. The bacteria evolved higher relative nitrogen and phosphorus content, consistent with selection for increased use of the more abundant elements. Total carbon assimilated also increased, indicating more efficient use of the limiting element. Altogether, our study shows that stoichiometry evolved over a relatively short time-period, and that it did so in a predictable direction given the carbon-limiting environment.

## Introduction

At the coarsest level, organisms consist of a mixture of elements in varying proportions. Common patterns in the ratios of elements are seen across life on Earth: all life requires large quantities of carbon, hydrogen, oxygen, nitrogen and phosphorus, and all life requires more carbon than nitrogen and more nitrogen than phosphorus. Because the cycling of carbon, nitrogen and phosphorus in ecosystems is driven in large part by biological processes, these elements tend to be the main focus of ecological stoichiometry (Sterner and Elser 2002). While all life shares broadly similar elemental profiles, the ratios vary substantially both inter-and intra-specifically (Vanni et al. 2002; Klausmeier et al. 2004; Bertram et al. 2008; Zimmerman et al. 2014). Over several decades, the field of ecological stoichiometry has established that stoichiometric variation has important ecological consequences. However, much less is known about the evolutionary origins of variation in organismal stoichiometry. It is not known whether organismal stoichiometry is a labile trait that can evolve fairly quickly when conditions change or if it is more constrained. Addressing this lack of knowledge is important because evolved changes in organismal stoichiometry might alter ecosystem processes (Elser 2006a) as well as shape responses to anthropogenic changes (Munday et al. 2013). In this paper, we present evidence from a laboratory experiment that changes in stoichiometry can evolve over an observable timescale and in predictable directions.

Variation in organismal stoichiometry affects many different levels of biological organization. At the physiological level, the growth-rate hypothesis predicts a correlation between the growth rate of individual organisms and their phosphorus content, driven by the greater need for phosphorus-rich ribosomes in rapidly growing organisms (Sterner and Elser 2002; Elser et al. 2003). Many groups of organisms exhibit this correlation, including zooplankton (Acharya et al. 2004), insects (Elser et al. 2006; González et al. 2014), phytoplankton (Hillebrand et al. 2013), and heterotrophic bacteria (Makino et al. 2003; Makino and Cotner 2004; Chrzanowski and Grover 2008). Differences in stoichiometry among organisms can also affect community and ecosystem processes such as predator-prey interactions (Meunier et al. 2012), competitive interactions (Gurung et al. 1999), and nutrient cycling (Elser et al. 1998; Vanni et al. 2002).

The relative availability of elements in the environment plays an important role in organismal stoichiometry. As a particular element becomes increasingly scarce, the proportion of that element in biomass generally declines, although the degree of physiological plasticity varies among organisms and across elements (Sterner and Elser 2002). However, the effect of elemental abundance on the evolution of organismal stoichiometry is not well established, however. All else being equal, one would expect selection for efficient nutrient use or “sparing”, such that the proportion of a scarce element in biomass would decrease over evolutionary time because selection favors organisms that require less of that element. However, the proportion of a scarce element in biomass might instead increase over evolutionary time. The adaptive value of that element for performing important functions could outweigh selection for nutrient sparing. Also, the evolution of improved mechanisms to acquire a scarce element could alleviate the effects of its scarcity. Because organisms often exhibit plasticity in response to nutrient scarcity, the improved uptake of a scarce element could lead to a higher proportion of that element in an organism’s biomass (Bragg and Wagner 2007).

Current evidence in support of the evolution of nutrient sparing is mixed. Several comparative studies of the elemental composition of proteins support the nutrient-sparing hypothesis. For example, in both *Escherichia coli* and yeast, proteins involved in the acquisition of nitrogen, phosphorus and sulfur each have a lower content of the element they acquire, suggesting that selection acted to reduce the nutrient content of proteins expressed under nutrient scarcity (Baudouin-Cornu et al. 2001). Terrestrial plants, whose growth is frequently limited by nitrogen, exhibit evidence of nitrogen sparing in their genomes and proteomes (Elser 2006b; Acquisti et al. 2009; but see Günther et al. 2013). Additionally, a recent study of a *Daphnia pulicaria* population, based on rearing individuals from an egg bank that had been deposited over many centuries in lake sediments, found that relaxation of phosphorus limitation in the lake led to reduced phosphorus-use efficiency in the *Daphnia* (Frisch et al. 2014). However, contrary to the predictions of the nutrient-sparing hypothesis, Bragg and Wagner (2007) found that proteins whose expression decreased in yeast experimentally evolved under carbon limitation were disproportionately carbon poor compared to the rest of the proteome. They suggested that the evolution of improved uptake of carbon might have alleviated the degree of carbon limitation, thereby allowing greater expression of carbon-rich proteins. The experiments analyzed by Bragg and Wagner (2007) ran for only a few hundred generations, so their results might reflect a short-term process whereas nutrient sparing might dominate over the long term.

The potential for evolved changes in stoichiometry has important implications for understanding how organisms will respond to anthropogenic change. The phytoplankton of the world’s oceans provide a critical ecosystem service by taking up atmospheric carbon dioxide. Half of all carbon removal from the atmosphere is performed by oceanic phytoplankton (Field et al. 1998), and some of this carbon is then exported to the ocean sediments, which act as a long-term carbon sink (Longhurst and Harrison 1989; Richardson and Jackson 2007). In order to make predictions about future climate change, it is therefore important to understand how phytoplankton growth will be affected by changing environmental conditions. A number of models (Quere et al. 2005; Follows et al. 2007) make such predictions based on our knowledge of phytoplankton ecology, but none of them incorporate evolutionary changes (Munday et al. 2013). Increases in sea-surface temperatures have already led to declines in phytoplankton biomass, associated with more severe nutrient limitation (Boyce et al. 2010). One process that might mitigate the effect of declining nutrient availability is evolution of changes in the stoichiometric ratios of elements in phytoplankton biomass. Evolution of stoichiometry could also affect the consequences of anthropogenic increases in nutrient availability in other ecosystems, such as those caused by runoff from agricultural systems into aquatic environments.

Many questions need to be answered before we can predict how stoichiometry might evolve in response to global change. Over what time spans can it evolve and at what rate? Does selection generally favor the reduced use of scarce resources? Is the proportional use of some elements more evolutionarily flexible than others? We can begin to address these questions using experimental evolution, in which organisms are allowed to evolve under controlled conditions in the laboratory. Microbes are valuable subjects for experimental evolution because they can evolve quickly due to their large population sizes and short generation times (Lenski et al. 1991). An additional benefit is that many microbes can be frozen throughout the course of the experiment and later revived for analysis. This feature allows ancestral and evolved organisms to be compared directly, under the same conditions in which they evolved.

To examine the potential for evolutionary change in organismal stoichiometry, we compared the carbon, nitrogen and phosphorus content of biomass in ancestral and evolved *E. coli* cells from a 50,000-generation laboratory evolution experiment (Lenski et al. 1991; Wiser et al. 2013). We predicted that the evolved bacteria would have lower carbon content, relative to nitrogen and phosphorus, compared to the ancestral strain for two reasons. First, the evolving bacteria have been maintained under carbon limitation in a medium that contains high concentrations of nitrogen and phosphorus (Lenski et al. 1991). All else being equal, we expected that such conditions should favor selection for reduced use of carbon in biomass, as predicted by the nutrient-sparing hypothesis. Similarly, the high concentrations of nitrogen and phosphorus should release the bacteria from any prior selection for reduced use of those elements. Second, the bacteria undergo daily batch transfer to fresh medium, a condition that selects for faster growth rates (Vasi et al. 1994). According to the growth-rate hypothesis (Sterner and Elser 2002), this regime should also select for an increase in the relative proportion of phosphorus in the bacterial biomass.

## Methods

### Long-term evolution experiment

The long-term evolution experiment (LTEE) was founded with *E. coli* B strain REL606 (Lenski et al. 1991; Jeong et al. 2009). Six of the 12 populations started directly from REL606; The other six populations began with a mutant clone, REL607, that differed by a neutral marker. The populations have been maintained in Davis-Mingioli minimal medium supplemented with 25 mg glucose and 2 mg thiamine per liter (DM25). The bacteria can use both glucose and thiamine as a carbon sources, but not citrate (another component of DM25 that the ancestral strain and most evolved populations cannot use). Given the glucose and thiamine in DM25, the molar ratios of C:N:P available to the bacteria are 1:17:50. For the one population, designated Ara-3, that evolved the ability to consume citrate (Blount et al. 2008; Blount et al. 2012), the available C:N:P ratios are 1.1:1:3. Each population is serially propagated by a 1:100 dilution into fresh medium each day; it then grows 100-fold until the available carbon is depleted, and it thus goes through ∼6.6 generations per day. Samples of each population are frozen every 500 generations. Portions of these frozen cultures can be revived, allowing direct comparison of ancestral and evolved bacteria. Lenski et al. (1991) provide more details about the methods used in the LTEE.

### Sample collection for stoichiometric analyses

We used a paired sampling design in which a clone isolated from each of the 12 populations at 50,000 generations was paired with its ancestral clone, either REL606 or REL607. Each clone was revived from a frozen stock and conditioned for one day in Luria-Bertani medium. After an additional day of conditioning in DM25, each clone was transferred, via a 1:100 dilution, to 500 mL of DM25. After 24 h, each culture was then split into two 250-mL aliquots, each of which was filtered through a pre-combusted glass-fiber filter. In the case of population Ara-3, which evolved the ability to consume citrate, we filtered only 125 mL due to the increased density of the population. Each filter was rinsed by filtering an additional 250 mL of 8.5% sterile saline solution made with distilled deionized water in order to remove dissolved nutrients present in the medium from the filter. The dry mass of each filter was measured before and after filtering. For each culture, one filter was analyzed for phosphorus content and the other for both carbon and nitrogen content.

We performed sample collection on four different dates with six cultures (three pairs of ancestral and evolved clones) filtered on each day. Pairs of ancestral and evolved clones were assigned to sampling dates randomly. On each day, 250 mL of sterile DM25 was filtered through each of two filters and processed identically to all other samples. These filters served as blanks in subsequent analyses.

To confirm that glass-fiber filters effectively collected both evolved and ancestral bacteria, we compared the number of colony-forming units in filtered and unfiltered medium. The filters retained more than 99% of both ancestral and evolved bacteria.

### Elemental analysis

Elemental content was measured by the Nutrient Analytical Services Laboratory at the University of Maryland Center for Environmental Science. Phosphorus mass was determined via extraction to dissolved phosphate followed by molybdenum blue colorimetric analysis (Aspila et al. 1976) using an Aquakem 250 photometric analyzer. Carbon and nitrogen content were measured by elemental analysis (U.S. EPA 1997) using an Exeter Analytical Model CE-440 Elemental Analyzer.

### Nucleic-acid content

The total nucleic-acid content of ancestral and evolved clones in stationary phase was measured after 24 h in DM25. The bacteria were revived from frozen stocks and conditioned as described above. Cells were then lysed by heating, and their nucleic-acid concentration was measured in duplicate using the Quant-iT RiboGreen assay kit.

### Data analyses

All values for biomass, carbon, nitrogen and phosphorus were corrected for background levels by subtracting the corresponding value measured from a blank filter on the same day. Background levels were always small (< 15%) relative to the measured value. Biomass was calculated as the change in dry mass of the filter before and after filtering. The percentages of carbon, nitrogen and phosphorus were calculated as the mass of each element divided by the total biomass. All nutrient ratios were calculated as molar ratios.

Two-tailed paired *t*-tests were used to test for differences between evolved and ancestral biomass, nucleic-acid content, percentages of C, N and P, and molar C:N, C:P and N:P ratios. Statistical analyses were conducted using JMP 9.0 software.

## Results

### Carbon, nitrogen and phosphorus content

We observed substantial differences in the ratios of carbon, nitrogen and phosphorus between the ancestral and evolved clones (fig. 1). Among the evolved clones, however, one had distinctly different C:N and C:P ratios compared to the other eleven, and that one was from the population that evolved the ability to consume the citrate present in the medium. As explained in the Methods, this population subsequently experienced conditions with much higher carbon availability than the other populations. The sample from this population reached a biomass concentration of 108.7 mg/L at 24 h, whereas the samples from the other populations had biomass concentrations between 10 and 18 mg/L. The citrate-consuming clone also had the highest measured C:N and C:P ratios of any of the evolved clones, with ratios as high as or higher than any samples of the ancestral clones. Given the distinctive growing conditions experienced by this clone, we excluded it from our statistical analyses. (Note, however, that inclusion of the citrate consuming clone does not affect whether the comparisons reported below are statistically significant at the 0.05 level, with two exceptions: the N:P ratio would no longer differ between the ancestral and evolved clones, while their nucleic-acid content per biomass would differ.)

**Figure 1:**
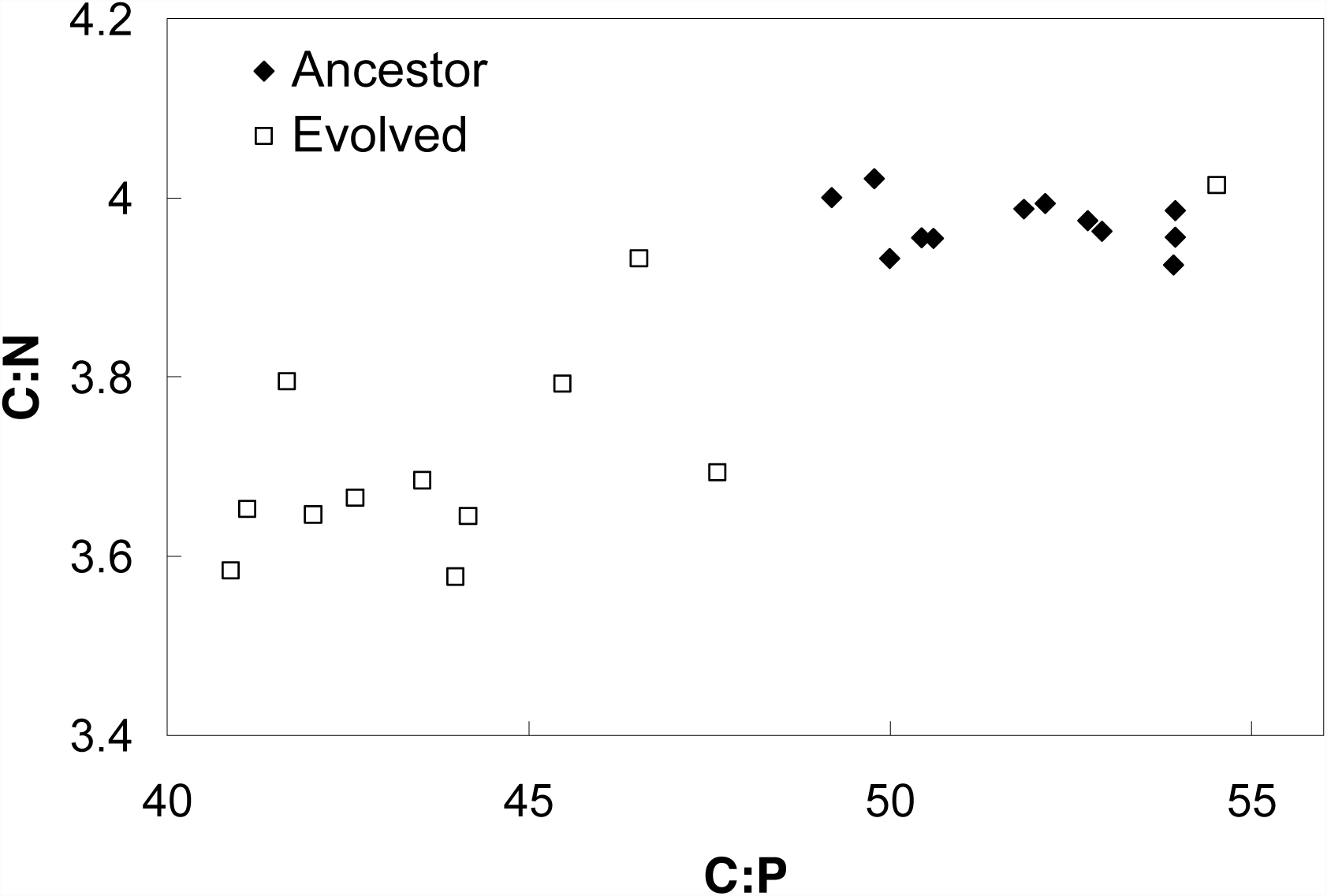
The molar C:N and C:P ratios of the ancestral (filled diamonds) and 50,000-generation evolved (open squares) clones from the long-term evolution experiment with *E. coli*. The evolved clone at the far upper right is from the population that evolved the ability to consume citrate, and it was excluded from the statistical analysis owing to the much higher carbon availability that it experienced.

The average molar C:N and C:P ratios both decreased significantly in the evolved clones compared to their ancestors (fig. 1, paired, two-tailed *t*-tests, C:N *P* < .001, C:P *P* < .001). Also, the average molar N:P ratio declined from 13.1 to 11.8 (*P* < .001). These changes in C:N and C:P ratios were driven by increases in the percentages of both nitrogen and phosphorus in the evolved clones. The percentage of biomass comprised of nitrogen atoms increased from 11.1% to 12.0% (*P* = .012), and that of phosphorus increased from 1.9% to 2.3% (*P* < .001). The percentage of carbon did not significantly change (fig. 2A, *P* = .888).

**Figure 2:**
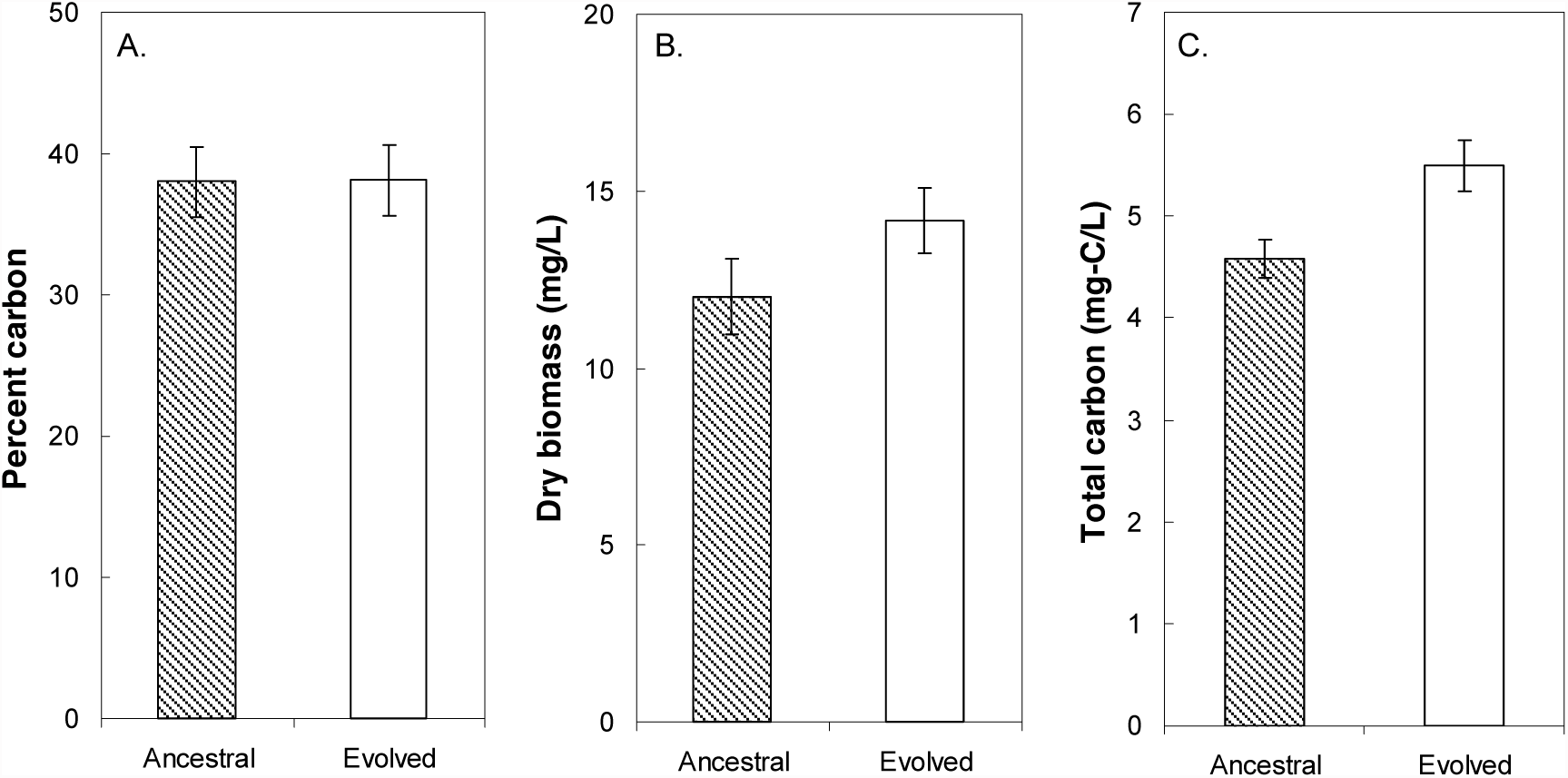
A. The percent carbon in cellular biomass did not differ between ancestral and evolved clones (*P* = 0.888). B. Total dry biomass was significantly higher in the evolved clones than in the ancestral clones (*P* < 0.001). C. The total carbon retained in biomass was also significantly higher in the evolved than the ancestral clones (*P* < 0.001). All data shown are means with 95% confidence intervals.

The significant increases in the percentages of both nitrogen and phosphorus in the evolved clones’ biomass implies that the proportion of one or more other elements must have declined. One possibility is a decline in the percentage of some unmeasured element, such as hydrogen, oxygen, sulfur or potassium. The other possibility is that the carbon content declined slightly, but that the decline was not detected. An evolved carbon content of 36.7% would be sufficient to offset the combined 1.3% increase in the nitrogen and phosphorus content; that level is well within the 95% confidence interval of 35.6 to 40.6% for the carbon content of the evolved bacteria.

### Biomass and carbon retention

The total bacterial biomass concentration increased by 17.7%, from 12.3 to 14.5 mg/L, in the evolved bacteria (Fig. 2B, *P* < 0.001). Although the percentage of carbon in the biomass did not change significantly, the total amount of carbon retained increased by 17.8%, from 386 to 455 μmol C/L (fig. 2C, *P* < .001), thus mirroring the increase in total biomass.

### Nucleic-acid content

The nucleic-acid content per culture volume increased 23.6%, from 344 to 425 μg/mL (*P* < .001), in the evolved bacteria. However, most of this change reflected the increase in overall biomass. The proportion of nucleic acid in the biomass averaged 28.3 μg-nucleic acid/mg-biomass in the ancestral clones and 29.6 μg-nucleic acid/mg-biomass in the evolved clones. This difference was not significant (*P* = .116), although the trend was in the predicted direction. Given that nucleic acids are about 8.7% P (Sterner and Elser 2002), then the change in nucleic-acid content could account for at most a 0.1 ug-p/mg-biomass increase in phosphorus content, which constitutes only ∼3% of the observed 3.6 ug-p/mg-biomass increase in phosphorus content per biomass.

## Discussion

Our results demonstrate that organismal stoichiometry can evolve quickly, at least in microorganisms, with changes observed over an experimental time span, albeit a long one that has run for decades (Fig. 1). To our knowledge, this is the first experimental observation of evolutionary changes in the overall stoichiometry of organisms, rather than inferred from observational evidence. The average C:P ratio decreased by 14% and the average C:N ratio declined by 6% during the evolution experiment. These changes were evolved, heritable changes that occurred in addition to any plastic physiological response. Previous work has shown that *E. coli* cells also exhibit a plastic response to variation in nutrient supply, with their C:P ratio decreasing ∼25% in response to declines in the C:P supply ratio, while the C:N supply ratio was held constant (Makino et al. 2003, C:N did not vary). Overall, these results indicate that evolutionary changes, even over an experimental time scale, can add to plastic physiological responses and further alter organismal stoichiometry. Moreover, these evolutionary changes can be of comparable magnitude to the plastic responses.

As predicted by the nutrient-sparing hypothesis, we observed significant declines in both the C:N and C:P ratios in biomass. These changes are consistent with selection for decreased carbon use relative to the use of nitrogen and phosphorus. However, we cannot distinguish the direct effect of selection due to carbon limitation from the indirect effects of selection favoring other traits in the evolution experiment. For example, some portion of the stoichiometric changes that we observed could be correlated responses to selection for larger cell size (Mongold and Lenski 1996), faster growth rate (Vasi et al. 1994), or other traits.

The uniquely high C:N and C:P ratios of the clone from the citrate-consuming lineage, which had access to 10 times as much carbon as any other population (Blount et al. 2008), provides further evidence that the declines in the C:N and C:P ratios in the other populations were beneficial specifically under the very low C:N and C:P supply ratios of the LTEE. However, the higher relative carbon content of the citrate-consuming clone is not necessarily an evolutionary response; its higher carbon content might also be, in whole or in part, a plastic physiological response to the higher carbon availability that it experiences in the experimental medium of the LTEE.

The proportion of cellular phosphorus contained in nucleic acids was much lower in our study than in previous experiments with *E. coli* (Makino et al. 2003). Our measurements were made during stationary phase, when carbon has been exhausted, rather than during exponential growth. Both DNA and RNA levels are closely tied to growth rate in bacteria, because the number of genome copies and transcription rate are higher in growing cells (Schaechter et al. 1958; Akerlund et al. 1995). Therefore, it is not surprising that nucleic-acid content would be quite low during stationary phase. The low nucleic-acid content, together with the lack of any significant change in the nucleic-acid content per biomass, indicates that the overall increase in the percentage of phosphorus in evolved cells reflects changes in cellular components other than nucleic acids. Based on the growth-rate hypothesis, we expected to see higher nucleic-acid content in the evolved cells because they exhibit a shorter lag time and faster growth rate upon exiting stationary phase (Vasi et al. 1994). However, our results are not a direct test of the growth-rate hypothesis, which focuses on growing organisms, rather than ones in stationary phase.

Given the carbon-limited conditions of the LTEE, one might reasonably expect that the strongest selection would be to reduce the proportion of carbon in the bacterial biomass. Therefore, it is notable that increases in nitrogen and phosphorus content drove the changes in the C:N and C:P ratios, while there was no significant change in the percentage of carbon in the biomass. Taken at face value, this finding suggests that the proportion of carbon in biomass may be less evolutionarily flexible than the proportions of nitrogen and phosphorus. Alternatively, changes in carbon content may be more difficult to detect because carbon makes up a much larger portion of biomass.

Although the proportion of carbon in biomass remained constant, the total amount of carbon that accumulated in the biomass increased significantly in the evolved bacteria (Fig. 2B). This increase was associated with an increase in the total biomass (Fig. 2C). The increase in carbon retention was not caused by an increase in the amount of carbon consumed because all of the bacteria (with the exception of the citrate consumer that was excluded from analysis) consumed the same 25 mg/L of glucose. Therefore, to retain more carbon in biomass, the bacteria must have released less carbon, either as carbon dioxide or byproducts excreted into the medium (and not then recycled for growth). Our results parallel those of an experiment in which yeast were evolved under carbon limitation for 350 generations (Goddard and Bradford 2003). The evolved yeast retained more carbon in biomass, but the proportion of carbon in biomass remained unchanged. It may be that carbon-use efficiency is more evolutionarily flexible than the proportion of carbon in biomass, at least in some organisms and contexts.

In this regard, it is also interesting that the LTEE provides no evidence of a tradeoff between growth rate and carbon-use efficiency. Instead, the evolved bacteria have increased in both their growth rate (Vasi et al. 1994; Novak et al. 2006) and carbon-use efficiency (this study; see also Vasi et al. 1994; Lenski and Mongold 2000; Novak et al. 2006 for indirect evidence based on optical density and total biovolume of cells, rather than on biomass). A tradeoff between growth rate and yield is one of the most well-established patterns in life history traits (Pfeiffer et al. 2001; MacLean 2008; Bachmann et al. 2013; Meyer et al. 2015). Under the unstructured (i.e., well mixed) conditions of the LTEE, selection for increased growth rate is strong and direct, whereas there is no direct selection for increased yield (Vasi et al. 1994); for example, a mutant that grows faster, despite using resources less efficiently, would still have a competitive advantage in the LTEE because the reduced resources would affect all competitors equally. Thus, it is surprising to observe an increase in biomass. It appears, therefore, that the ancestor was sufficiently maladapted to the conditions of the LTEE that both growth rate and yield could increase (Novak et al. 2006).

The LTEE is an ongoing experiment, and the trend of decreasing C:N and C:P ratios might well continue over longer time spans. The most likely mechanisms for the changes in stoichiometry observed thus far are changes in the abundance of proteins and other broad classes of cellular components that differ in their elemental content. Over longer timescales, selection for nutrient sparing might also change the composition of individual proteins or other specific components. We focus on proteins in the discussion below because we can easily calculate the effect of individual mutations on the elemental content of proteins. However, similar logic applies for other cellular components.

Comparative studies have found that selection can affect the amino-acid content of proteins in organisms, reducing their use of the most limiting elements (Baudouin-Cornu et al. 2001; Elser 2006b). However, the fitness benefit of any mutation changing a single amino acid is likely to be extremely small. Following the approach of Bragg and Wagner (2009), and given the effective population size for the LTEE of 3.3 x 10^7^ (Lenski et al. 1991), we calculate that mutations that save a single carbon atom would be visible to selection, such that *s* > 1/(2*N*_e_) (Kimura 1983), in proteins with more than ∼450 copies per cell. Individual amino acid changes, specifically Trp to Gly, can save as many as 9 carbon atoms per protein molecule (Bragg and Wagner 2009). A mutation that saved 9 carbon atoms per protein molecule could be selected for in proteins with ∼50 or more copies per cell. Of course, such a mutation would require a very long time to achieve fixation. First, the mutation would have to escape drift loss while it was still rare; a successful escape would require many millions of mutational events, given that the probability of escaping drift loss is only ∼4*s* (Gerrish and Lenski 1998). Also, in the asexual regime of the LTEE, large-effect mutations are expected to drive out smaller effect mutations due to clonal interference (Gerrish and Lenski 1998). For example, consider a mutation that saves 9 carbon atoms per molecule in an abundant protein with 10,000 copies per cell. Even after escaping drift and without competition from other mutations, this mutation would require on the order of 10^6^ generations (∼400 years) to approach fixation in the population.

Our results demonstrate the potential for experimental studies to shed new light on evolutionary changes in stoichiometry, since measurable inherited changes can occur over experimentally tractable time scales. The relatively rapid evolutionary changes in stoichiometry also raise the possibility that large-scale changes in nutrient availability in Earth’s ecosystems, for example those caused by climate change or increased fertilizer use, could drive evolutionary changes in organismal stoichiometry. Shifts in organismal stoichiometry could, in turn, alter the effects of anthropogenic change on ecological interactions and ecosystem processes. Evolution of stoichiometry might be usefully incorporated into eco-evolutionary models, such as those that forecast the response of global phytoplankton communities to climate change (Follows et al. 2007; Munday et al. 2013). In order to successfully model the evolution of stoichiometry, however, we need to know much more about how stoichiometry responds to selection, and experimental evolution offers a valuable tool for acquiring that knowledge.

## Acknowledgments

We thank Neerja Hajela for assistance in the lab; Mike Wiser and Rohan Maddamsetti for comments on this paper; and Jim Elser and Marcia Kyle for preliminary nutrient analysis. This research was supported, in part, by an EPA STAR Fellowship to C.T., an NSF grant (DEB-1019989) to R.E.L., and the BEACON Center for the Study of Evolution in Action (NSF Cooperative Agreement DBI-0939454).

